# Single-cell whole-genome sequencing reveals mutational landscapes of DNA mismatch repair deficiency in mouse primary fibroblasts

**DOI:** 10.1101/668467

**Authors:** Lei Zhang, Xiao Dong, Xiaoxiao Hao, Moonsook Lee, Zhongxuan Chi, Bo Jin, Alexander Y. Maslov, Winfried Edelmann, Jan Vijg

## Abstract

DNA Mismatch repair (MMR) deficiency is a major cause of hereditary non-polyposis colorectal cancer, and is also associated with increased risk of several other cancers. This is generally ascribed to the role of MMR in avoiding mutations by correcting DNA replication errors. In MMR knockout mice very high frequencies of somatic mutations, up until 100-fold of background, have been reported. However, these results have been obtained using bacterial reporter transgenes, which are not representative for the genome overall, and mutational patterns of MMR deficiency remain largely unknown. To fill this knowledge gap, we performed single-cell whole-genome sequencing of lung fibroblasts of *Msh2*^−/−^ and wild-type mice. We observed a 4-fold increase of somatic single nucleotide variants (SNVs) in the fibroblasts of *Msh2*^−/−^ mice compared to those of wild-type mice. The SNV signature of *Msh2* deficiency was found to be driven by C>T and T>C transitions. By comparing it to human cancer signatures, we not only confirmed the inferred MMR-deficiency-related etiology of several cancer signatures but also suggested that MMR deficiency is likely the cause of a cancer signature with its etiology previously unknown. We also observed a 7-fold increase of somatic small insertions and deletions (INDELs) in the *Msh2*^−/−^ mice. An elevated INDEL frequency has also been found in human MMR-related cancers. INDELs and SNVs distributed differently across genomic features in the *Msh2*^−/−^ and control cells, with evidence of selection pressure and repair preference. These results provide insights into the landscape of somatic mutations in normal somatic cells caused by MMR deficiency.

**Significance:** Our results show that MMR deficiency in the mouse is associated with a much lower elevation of somatic mutation rates than previously reported and provides the first MMR whole-genome mutational landscapes in normal somatic cells *in vivo*.

## Introduction

Cancer is a genetic disease ultimately resulting from somatic mutations in normal cells driving phenotypes such as abnormal growth, metastasis and tissue invasion. It is, therefore, not surprising that an increase in spontaneous mutation rate accelerates cancer risk (1). A critical pathway that greatly increases mutation rate in normal cells when inactivated is DNA mismatch repair (MMR), which normally corrects base mismatches and insertion/deletion mispairs during DNA replication (2), thereby preventing such errors from turning into mutations. In humans, mutations in genes that control MMR have been shown to greatly increase the risk of various cancers, most notably Hereditary Nonpolyposis Colorectal Cancer (HNPCC) (3,4); in mice, MMR deficiency mostly increases lymphomas (5). Using selectable transgenic reporter systems it has previously been demonstrated that somatic mutation frequency in tissues from MMR knockout mice greatly exceeded that in normal control mice (6). For example, using the reporter gene *supFG1*, extremely high mutation frequencies were observed in the normal skin and colon of MMR-deficient mice (6). However, reporter genes are not necessarily representative for the genome overall and suffer from potential artifacts (7).

Due to their low abundance, *de novo* mutations in normal tissues cannot be directly detected except in clonal lineages such as cancers. To obtain an accurate and comprehensive insight into the mutational landscape of normal cells and tissues single-cell approaches are needed (8). We recently developed an accurate single-cell whole-genome sequencing assay for quantitatively analyzing *de novo* somatic mutations. This method, single-cell multiple displacement amplification (SCMDA), was validated by comparing mutation frequency and spectrum in whole-genome amplified cells with those in clones of fibroblasts from the same cell population without amplification (9). In this present study, we applied SCMDA to study single lung fibroblasts of *Msh2*^−/−^ mice as compared to their wild-type equivalents, i.e., cells from *Msh2*^+/+^ littermate controls as well as non-littermate wild-type controls. As expected, a significantly elevated frequency of both somatic single-nucleotide variations (SNVs) and somatic small insertions and deletions (INDELs) in *Msh2*^−/−^ cells was observed, albeit orders of magnitude less than previously reported using reporter transgenes in Msh2 knockout mice. Furthermore, we documented the mutation signature of *Msh2*-deficiency in these normal somatic cells in relation to the signatures previously observed in human cancers and analyzed the distribution of mutations across genomic features.

## Materials and Methods

Detailed methods are provided in Supplementary Materials and Methods.

### Transgenic mice and genotyping

Mice nullizygous for the *Msh2* gene, were generated and backcrossed into C57BL/6 as described previously (10). Genotypes were validated by PCR of tail DNA (Fig. S1). In this study, three *Msh2*^−/−^ mice (4-5 months), two of their wild-type littermates (4-5 months), and two additional, non-littermate wild-type mice (C57BL/6, 6 months) were used. All procedures involving animals were approved by the Institutional Animal Care and Use Committee (IACUC) of Albert Einstein College of Medicine and performed in accordance with relevant guidelines and regulations.

### Single lung fibroblast isolation

Primary lung fibroblasts were isolated following a cell isolation protocol adapted from Seluanov *et al* (11). In brief, mouse lung was minced and incubated in DMEM F-12 medium with 0.13 unit/ml Liberase Blendzyme 3 and 1x penicillin/streptomycin at 37°C for 40 min. Dissociated cells were washed, plated in cell culture dishes with complete DMEM F-12 medium, 15% FBS and cultured at 37°C, 5% CO_2_, 3% O_2_. When reaching confluence, cells were split and replated in EMEM medium supplemented with 15% FBS and 100 units/ml penicillin and streptomycin. Lung fibroblasts were purified by further passaging in the same medium.

Single lung fibroblasts were isolated using the CellRaft AIR system (Cell Microsystems) according to the manufacturer’s instructions. Isolated single fibroblasts in 2.5 μl PBS were frozen immediately on dry ice and kept at −80°C until amplification.

### Single-cell whole-genome amplification, library preparation and sequencing

The isolated single fibroblasts were amplified using SCMDA as described (9). The amplicons were subjected to quality control using a locus dropout test (12). Of those passing the quality control, three amplicons per mouse were subjected to library preparation and sequencing with 150-bp paired-end reads on an Illumina HiSeq X Ten sequencer (Novogene, Inc). The average sequencing depth across the genome was 27.5±3.2 (avg. ± s.d.), covering on average 86.9% of the genome of the single fibroblasts (Table S1). Bulk DNAs extracted from tails of the same mice were sequenced without amplification and used for filtering out germline polymorphisms during variant calling as described (9).

### Computational analyses

All methods of data analysis and statistics are provided in Supplementary Information.

### Data availability

All raw sequencing data will be available in SRA database upon publication.

## Results

### SNV frequency, spectrum and signature of Msh2^−/−^ mouse fibroblasts

In *Msh2*^−/−^ mouse fibroblasts we observed a SNV frequency of 5,343±2,450 (avg. ± s.d.) per cell after adjusting for genome coverage (Table S2). This frequency is approximately 4-fold higher than the mutation frequency observed in the same cell type of the wild-type mice (i.e., 1,382±394 per cell, *P*=0.0012, two-tailed *t* test, Fig. 1A), which is in the same range as what we previously observed in dermal fibroblasts of wild-type C57BL/6 mice (12). In addition to the elevated SNV frequency, *Msh2*^−/−^ mice displayed a trend of elevated mouse-to-mouse variance of SNV frequency as compared to the wild-type mice (Fig. 1A). Comparing the mutation spectra of the fibroblasts, we observed increased fractions of transition mutations, i.e., T>C and C>T, and decreased fractions of transversions in the *Msh2*^−/−^ cells as compared to the wild-type spectra (Fig. 1B).

**Figure 1.**
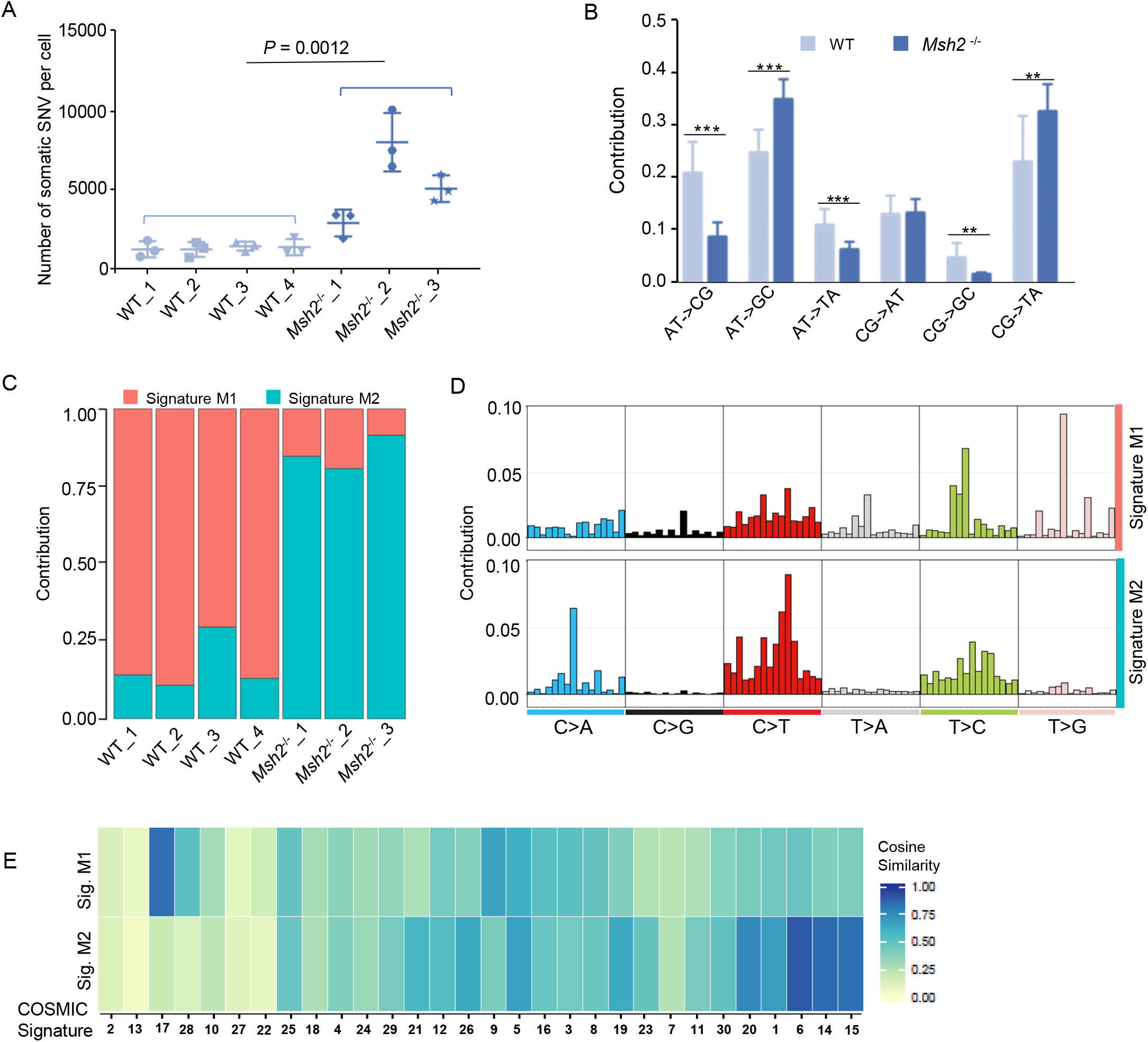
SNV frequency, spectrum and mutational signature of *Msh2*^−/−^ and wild-type (WT) mouse lung fibroblasts. **A.** Number of SNVs per cell. Error bars indicate standard deviations (s.d.). **B.** Spectrum of somatic SNVs. *P* value was presented by asterisks (***, *P* < 0.001; **, *P* < 0.01, two-tailed *t* test). **C.** Contributions of signatures M1 and M2 to total SNVs. **D.** Mutation signatures in the context of their flanking base pairs (in alphabetical order, e.g., in the first column C>A category from left to right, ApCpA > ApApA to TpCpT > TpApT). **E.** Cosine similarity (shown by heatmap) between signatures M1 and M2, and COSMIC signatures. The COSMIC signatures have been ordered according to hierarchical clustering.

To further investigate the role of *Msh2*-deficiency on mutational processes, we decomposed the mutation spectra into mutational signatures using non-negative matrix factorization (NMF) (13). To make the signatures comparable with human cancer studies, we humanized the mouse spectra before NMF, based on the difference of trinucleotide compositions between mouse genome and human exome according to ref. (14) (Fig. S2). Human exome was used because most studies of somatic mutations in tumors were based on whole-exome sequencing (15). Using NMF we identified two mutational signatures: M1 and M2 (Figs. 1C, D). As signature M2 substantially increased in the *Msh2*^−/−^ cells compared to wild-type, we considered M2 as the mutation signature of *Msh2*-deficiency in normal mouse lung fibroblasts.

In signature M2, C>T transitions occurred with increased frequencies of G<u>C</u>N and N<u>C</u>G trinucleotide contexts (N represents any one of the A, T, G and C bases) than other contexts. In comparison with human cancer signatures from the COSMIC study (15), signature M2 exhibits the highest similarity to COSMIC signature 6 (cosine similarity=0.902, Fig. 1E; Fig S3 and Table S3), which is associated with MMR-deficiency in cancers (15). Signature M2 is also highly correlated with three other MMR-deficiency-related COSMIC signatures, i.e., signatures 15, 20 and 26 (Fig. 1E; Fig S3 and Table S3). Of note, all the four MMR-deficiency-related COSMIC signatures in tumors are associated with high INDEL frequencies, which we also observed in the *Msh2*^−/−^ (see the following section). In addition to the correlation with the MMR-related COSMIC signatures, signature M2 showed the second highest correlation with a signature of unknown etiology, COSMIC signature 14 (cosine similarity=0.853). This suggests that COSMIC signature 14 is likely also a result of MMR deficiency. COSMIC signature 14 has been observed in uterine cancers and adult glioma samples with remarkably high SNV frequencies (> 200 mutations per Mb) (15).

### INDEL frequency, spectrum and genomic distribution

We found a 7-fold increase of INDEL frequency in *Msh2*^−/−^ mouse fibroblasts (3172.2±970.5 per cell) as compared to wild-type fibroblasts (426.3±155.5, *P*=2.43×10^−5^, Fig. 2A, Table S2). This increase is substantially higher than the increase of SNVs, i.e., 4-fold, resulting in a higher INDEL-to-SNV ratio in *Msh2*^−/−^ than in wild-type cells (0.59 and 0.31, respectively). As to the proportions of insertions and deletions, we found the majority (80.2%) in *Msh2*^−/−^ fibroblasts were deletions, while in wild-type fibroblasts most were insertions (77.2%; Fig. 2B).

**Figure 2.**
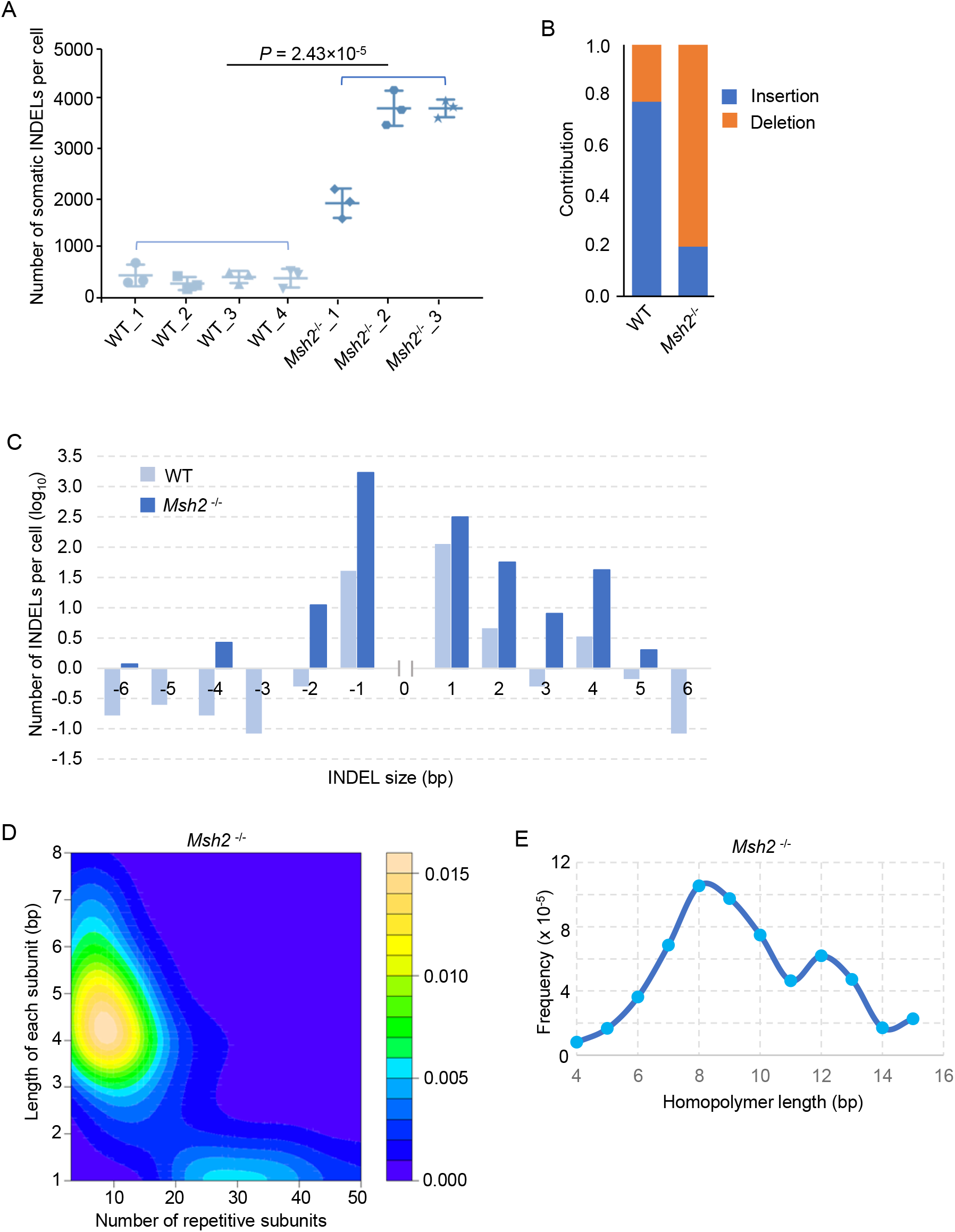
INDEL frequency, spectrum and distribution of *Msh2*^−/−^ and wild-type mouse lung fibroblasts. **A.** Number of INDELs per cell. Error bars indicate s.d.. **B.** Average contributions of insertions or deletions per cell. **C.** Size distribution of INDELs. Negative x-axis values present deletions and the positive present insertions. Only INDELs ≤ 6 bp are shown (Table S4 for all). **D.** Distribution of 1-bp INDELs in simple repeats in *Msh2*^−/−^ mice. Color bar indicates density. See Fig. S4A for wild-type mice. **E.** Frequency of 1-bp INDELs in homopolymers in *Msh2*^−/−^ mice. See Fig. S4B for wild-type mice.

Most of the INDELs of *Msh2*^−/−^ cells were 1-bp in length (Fig. 2C, Table S4). We analyzed the distribution of 1-bp INDELs in simple repeats and homopolymers (polymers composed of multiple identical mononucleotides), which are the most abundant classes of microsatellites in the human genome related to many human diseases (16,17). For simple repeats, the 1-bp INDELs preferably occurred at repeat regions with 5-8 repeat subunits of 4-5 bp subunit length (Fig. 2D). A similar distribution of INDELs in simple repeats was observed in the wild-type cells (Fig. S4A). For homopolymers, we found that the frequency of 1-bp-INDELs at a homopolymer length of 8 base pairs was the highest in both *Msh2*^−/−^ and wild-type cells (Figs. 2E and S4B-D), which may be due to preference of replicative polymerase slippage at homopolymers.

### SNV or INDEL distribution across genomic features

We analyzed distributions of SNVs and INDELs across different genomic features. In contrast to a marginal SNV depletion in introns, SNVs were significantly depleted in exons and UTRs, i.e., their occurrence was less than expected by chance alone in both wild-type and *Msh2*^−/−^ mice (Fig. 3A). The depletion in exons was less dramatic in cells of the *Msh2*^−/−^ mice than those of the wild-type mice (61.9% and 42.4%, respectively, of what would be expected by chance alone; Figs. 3A, B). This suggests that in wild-type mice, the major cause of depletion is a possibly higher MMR efficiency in exons than other genomic regions, and the effect of negative selection may be secondary.

**Figure 3.**
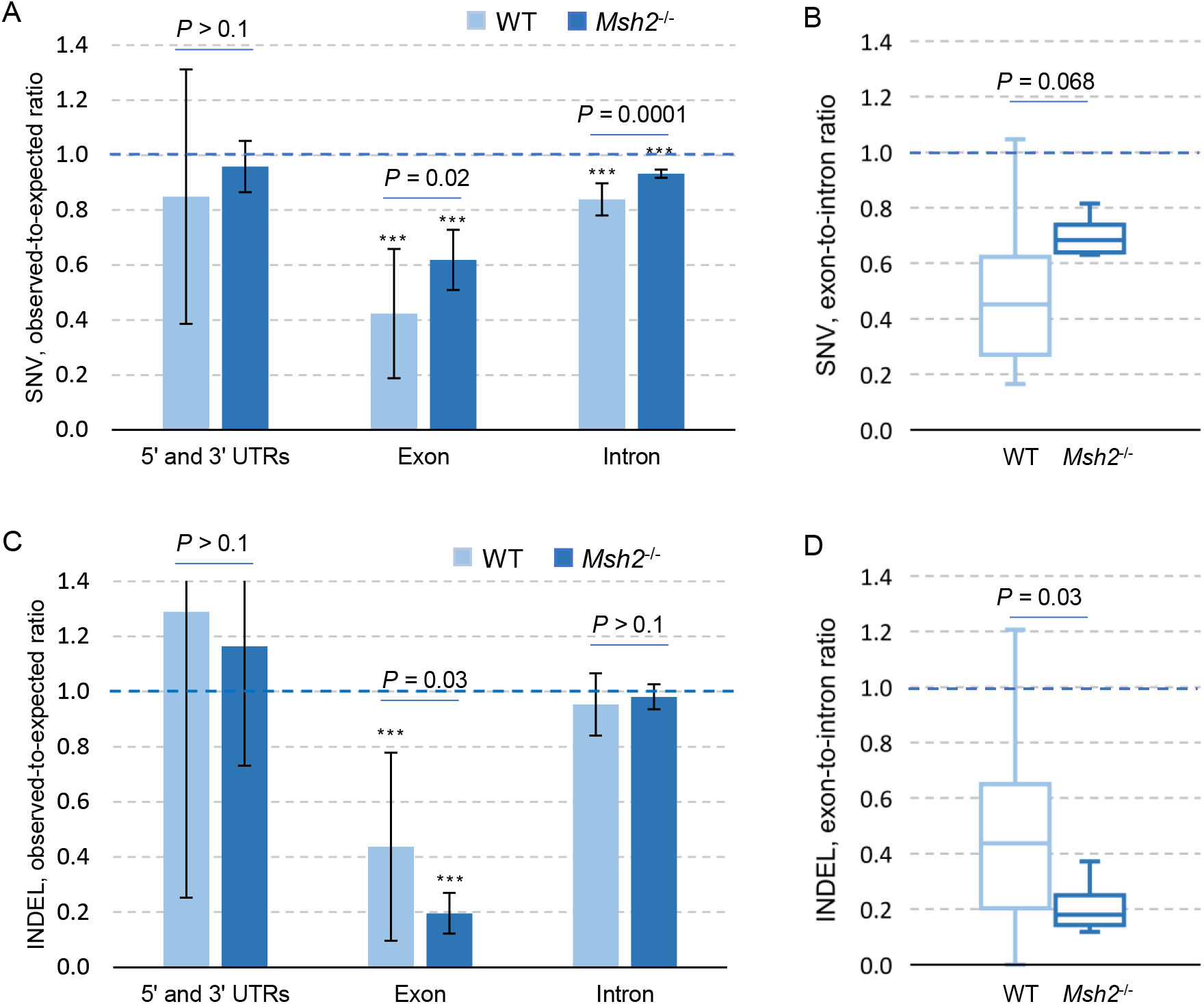
Distribution of SNVs and INDELs across the genomic features of *Msh2*^−/−^ and wild-type mouse lung fibroblasts. **A.** Ratio of observed number of SNVs to the number expected by chance alone. Bar height indicates the mean and error bar indicates s.d.. The statistical difference was tested by two-tailed *t* test. *P* values presented by asterisks indicate difference between the observed and the expected (***, *P* < 0.001; two-tailed *t* test). **B.** Ratio of SNV frequency in exons to that in introns per cell. **C.** Ratio of observed number of INDELs to the number expected by chance alone. *P* values presented by asterisks indicate difference between the observed and the expected. **D.** Ratio of INDEL frequency in exons to that in introns per cell.

For INDELs, in contrast to SNVs, a more dramatic depletion in the exome was observed in *Msh2*^−/−^ than in wild-type cells (19.6% and 43.7%, respectively, of what would be expected by chance alone; Figs. 3C, D and Table S5). Considering that exonic INDELs are much more likely to cause gene inactivation than exonic SNVs, these findings suggest that negative selection is the main cause of INDEL depletion from the exome.

## Discussion

The results of this study were based on utilization of single-cell whole-genome sequencing (9). This is the most accurate strategy for evaluation of *in vivo* mutation frequencies and spectra currently available (18). The 4-fold and 7-fold increase observed in SNVs and INDELs, respectively, is much more modest than the 50-fold increase for MSH2-deficiency previously reported using reporter mice, i.e., 35.7 to 555.1 × 10^−5^ mutation per base pair (equivalate to 2 to 31 × 10^6^ mutations per cell with diploid genome) in *Msh2*^−/−^ mice (6). These high numbers could be the result of artifacts, possibly associated with a lack of representation of a single bacterial reporter gene for the overall mouse genome. Although as shown recently, analysis can be conducted using single-cell clones or organoids in human cell lines (19), this strategy has the disadvantage of measuring mutations in progeny of stem or progenitor cells, in which mutation frequency is likely to be much lower than in their differentiated counterparts (20). Also, such strategies do not account for mutations arising during clone formation. Indeed, only single-cell sequencing allows direct mutational analysis of all types of cells even in postmitotic cells, e.g., neurons and muscle fibers (20–22).

An interesting question is when the mutations observed arose. SNVs are mostly a result of replication errors and since in adult organisms there is generally not much cell division except in intestine and the hematopoietic system, most somatic mutations observed in this study likely occurred during development. However, since we did not observe a significant number of mutations shared by different cells, selection against mutant embryonic cells likely prevents transmission of highly mutated cells to offspring. While the numbers of mutations we observed, i.e., about 5,000 per cell on average, are obviously compatible with survival, it is likely the maximum tolerated mutation load of this cell type *in vivo*. Indeed, *Msh2*^−/−^ mice live very short and die from lymphoma, which does not allow longitudinal studies to settle this issue. It is possible that the increased mutation loads of somatic cells leads to premature aging which has not been systematically studied for MMR-deficient mice.

*Msh2* deficiency was found to significantly affect the mutational spectra in normal cells. We found more transitions than transversions for SNVs and more deletions than insertions for INDELs in *Msh2*^−/−^ cells compared to wild-type cells. Similar results have been observed in skin or colon of *Msh2*^−/−^ mice with the reporter systems (6). However, we found a much smaller INDEL-to-SNV ratio than what has been observed using the reporter systems: 0.59 in our study compared to 1.58 and 29 with reporter genes *supFG1* and *cll,* respectively. Studies of somatic mutation spectra in MMR-related human cancers have been focused on the SNV signatures (15). By comparing SNV signatures of *Msh2*^−/−^ cells with COSMIC signatures, our results confirmed the inferred etiology of COSMIC signatures 6, 15, 20 and 26 as MMR deficiency. Importantly, our results also suggest a new causal link between MMR deficiency and cancer signatures with unknown etiology, i.e., COSMIC signature 14. This would be in keeping with a recent study reporting mutation signatures of tumors with concurrent loss of polymerase proofreading and mismatch repair function closely resembling the COSMIC signature 14 (23). Taken together, these results strongly suggest that COSMIC signature 14 is associated to and most likely a result of MMR deficiency.

We found that somatic SNVs and INDELs distributed differently across genomic features, especially exons. Depletion of SNVs from exons was greater in wild-type cells than in *Msh2*^−/−^ cells, which seems counterintuitive. However, this is likely a result of higher MMR efficiency in exons than in other genomic regions in the wild type, which is in agreement with previous findings of SNV distribution in human MMR-deficient tumors (24) and may be a result of H3K36me3-mediated MMR protecting exonic regions preferentially (25,26). In contrast to SNVs, INDELs in wild-type cells showed significant depletion in the exome, and the depletion was more dramatic in the *Msh2*^−/−^. Unlike SNVs, the majority of INDELs in exons causes reading-frame shifts, and subsequent inactivation of the gene’s activity. This suggests that INDEL depletion from the exome is due to negative selection in the tissue *in vivo* (27).

In summary, we directly identified *de novo* somatic SNVs and INDELs in primary cells from wild-type and *Msh2*^−/−^ mice, determined their mutational frequency, signatures as well as distribution across genomic features. The same strategy could be applied to studying normal cells in tissues during early development of tumors or even before cancers arise in their normal somatic progenitor cells.

## Authors’ Contributions

*Conception and design*: J. Vijg

*Development of methodology*: L. Zhang, X. Dong

*Acquisition of data (provided animals, acquired and managed patients, provided facilities, etc.)*: B. Jin, W. Edelmann

*Analysis and interpretation of data (e.g., statistical analysis, biostatistics, computational analysis)*: L.Zhang, X. Dong, X. Hao

*Writing, review, and/or revision of the manuscript*: L. Zhang, X. Dong, W. Edelmann, J. Vijg

*Administrative, technical, or material support (i.e., reporting or organizing data, constructing databases)*: X. Hao, M. Lee, Z. Chi, A.Y. Maslov

*Study supervision*: J. Vijg

## Acknowledgement

This project was supported by NIH grants P01 AG017242 (J. Vijg), P01 AG047200 (J. Vijg), U01 ES029519 (J. Vijg) and K99 AG056656 (X. Dong).

