## Supplementary Information for "Single-cell whole-genome sequencing reveals mutational landscapes of DNA mismatch repair deficiency in mouse primary fibroblasts"

#### DNA extraction and PCR genotyping

We extracted genomic DNA from tail of each mouse using the DNeasy Blood & Tissue Kit (Qiagen) following the manufacturer's specifications. The concentrations of DNA were quantified using the Qubit High Sensitivity dsDNA Kit (Invitrogen Life Science) and the qualities of DNAs were evaluated with 1% agarose gel electrophoresis. We performed PCR genotyping using the genomic DNA as PCR template. Each reaction contains 1 µl of gDNA (10ng/µl), 1.5 µl of 10x PCR buffer II (Roche), 1.5 µl of $MgCl_2$ (25mM, Roche), 0.1 µl of Taq Gold (5U/ µl) and Primer A, B and C (The sequences of Primers are listed in Fig. S1). The total reaction volume of PCR is 12.5 µl. PCR conditions were 94 °C for 5 min; and 40 cycles 94 °C for 45 s, 55 °C for 1 min and 72 °C for 1 min; and 72 °C for 5 min. The PCR results were shown in the picture of 1% agarose gel electrophoresis (Fig. S1).

#### Sequence alignment

Raw sequence reads were subject to quality control using FastQC (https://www.bioinformatics.babraham.ac.uk/projects/fastqc/), adaptor- and quality-trimmed using Trim Galore (https://www.bioinformatics.babraham.ac.uk/projects/trim_galore/), and aligned to reference genome mouse mm10 using bwa mem (1). PCR duplicates were removed using samtools (2). The aligned reads were then INDEL-realigned and base-pair score quality recalibrated using GATK (3).

#### Variant calling

We used the overlap of three variant callers as described (4). Somatic mutations were determined as the intersection of variant calls of three software tools, i.e., VarScan2 (5), Haplotypecaller (3) and MuTect2 (6), from a single cell using corresponding bulk as control with default parameters and a minimum depth of 20x in both single cell and bulk. Candidates were further filtered out if reported previously in dbSNP or had any variant-supporting reads in the bulk.

#### Analysis of mutation signatures

Somatic SNVs of 3 cells of each mouse were pooled together to represent the mutation spectrum of each mouse. We converted mouse spectrum to humanized spectrum by adjusting with the ratio of trinucleotide contexts in mouse genome to that of the human exome, using an R script provided in ref (7). We determined mutation signatures from humanized spectrum using non-negative matrix factorization (NMF) using R package "MutationalPatterns" (8).

### Annotation of variants with genome features

Gene annotation of mutations were performed using ANNOVAR (9) using mm10. Repeat region of the reference genome was downloaded from RepeatMasker (http://repeatmasker.org). Mutations in repeat regions were annotated using bedtools (10).

### Statistics

Statistical differences between wild-type and *Msh2*$^{-/-}$ mouse cells were determined by the unpaired *t*-test. Statistical significance was determined as *P*-value. The details of statistical analysis in each figure were described in the figure legends.
