## Supplementary Figures for "Single-cell whole-genome sequencing reveals mutational landscapes of DNA mismatch repair deficiency in mouse primary fibroblasts"

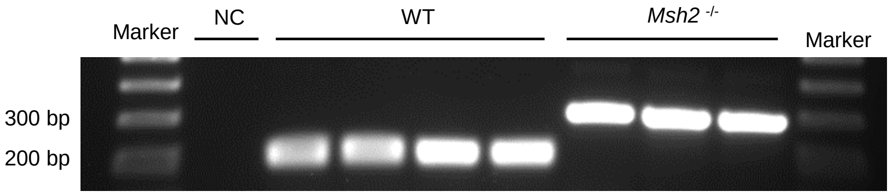


Primer A: CCCTCCTGTTGAGCCATCTTA

Primer B: GCCAGCTCATTCCTCCACTC

Primer C: TTCGCTGCTTGTCTCTGGAAT

**Fig. S1. PCR genotyping.**
In wild-type mice, the pair A/C was amplified; and in *Msh2*$^{-/-}$ the  pair A/B was amplified..

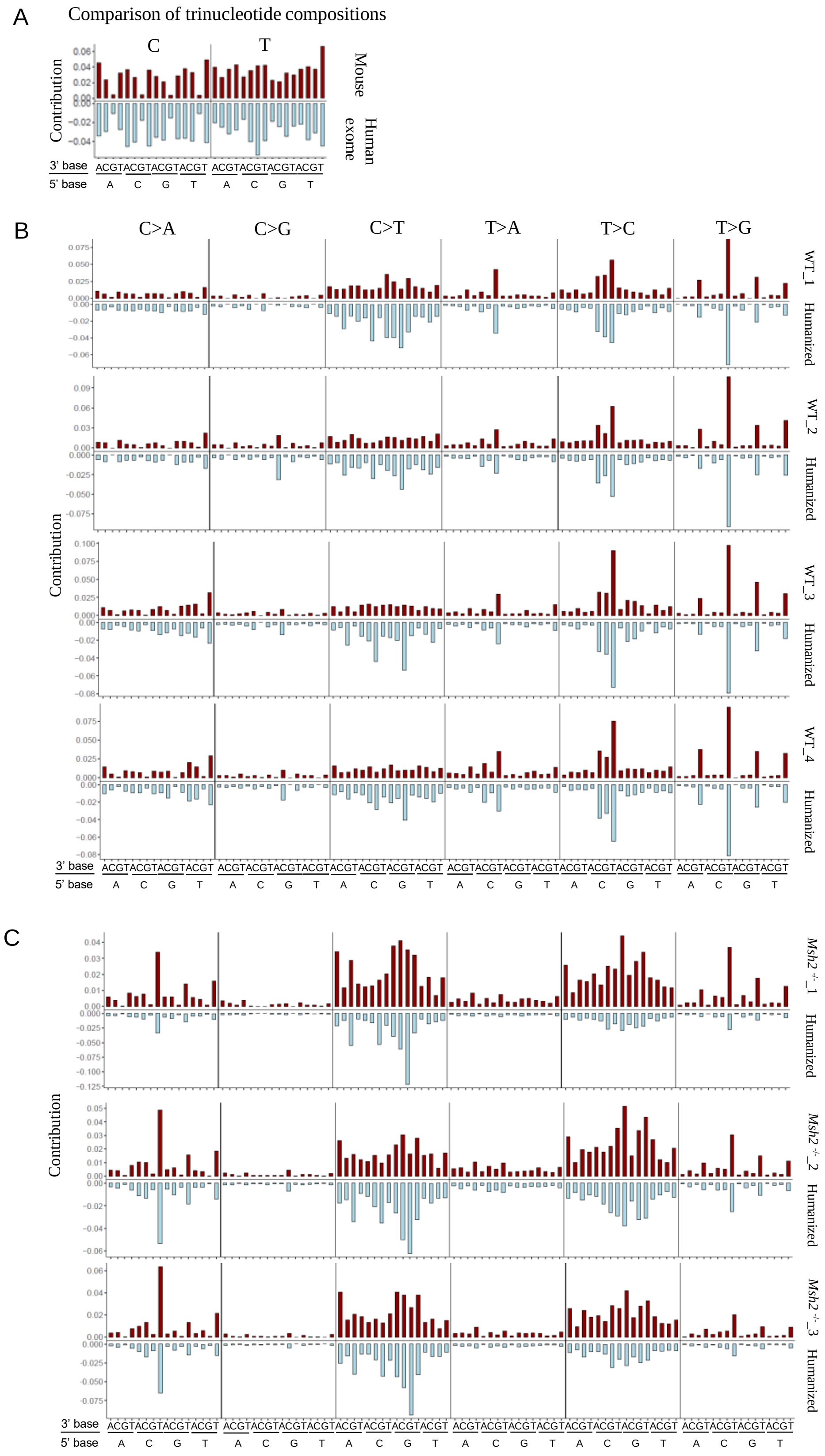


**Fig. S2. Mutation spectra and their corresponding humanized versions**. **A.** Contribution of trinucleotides in the mouse reference genome and the human exome. **B.** Contribution of trinucleotides in wild-type mice and their humanized versions. **C.** Contribution of trinucleotides in *Msh2*$^{-/-}$ mice and their humanized versions.

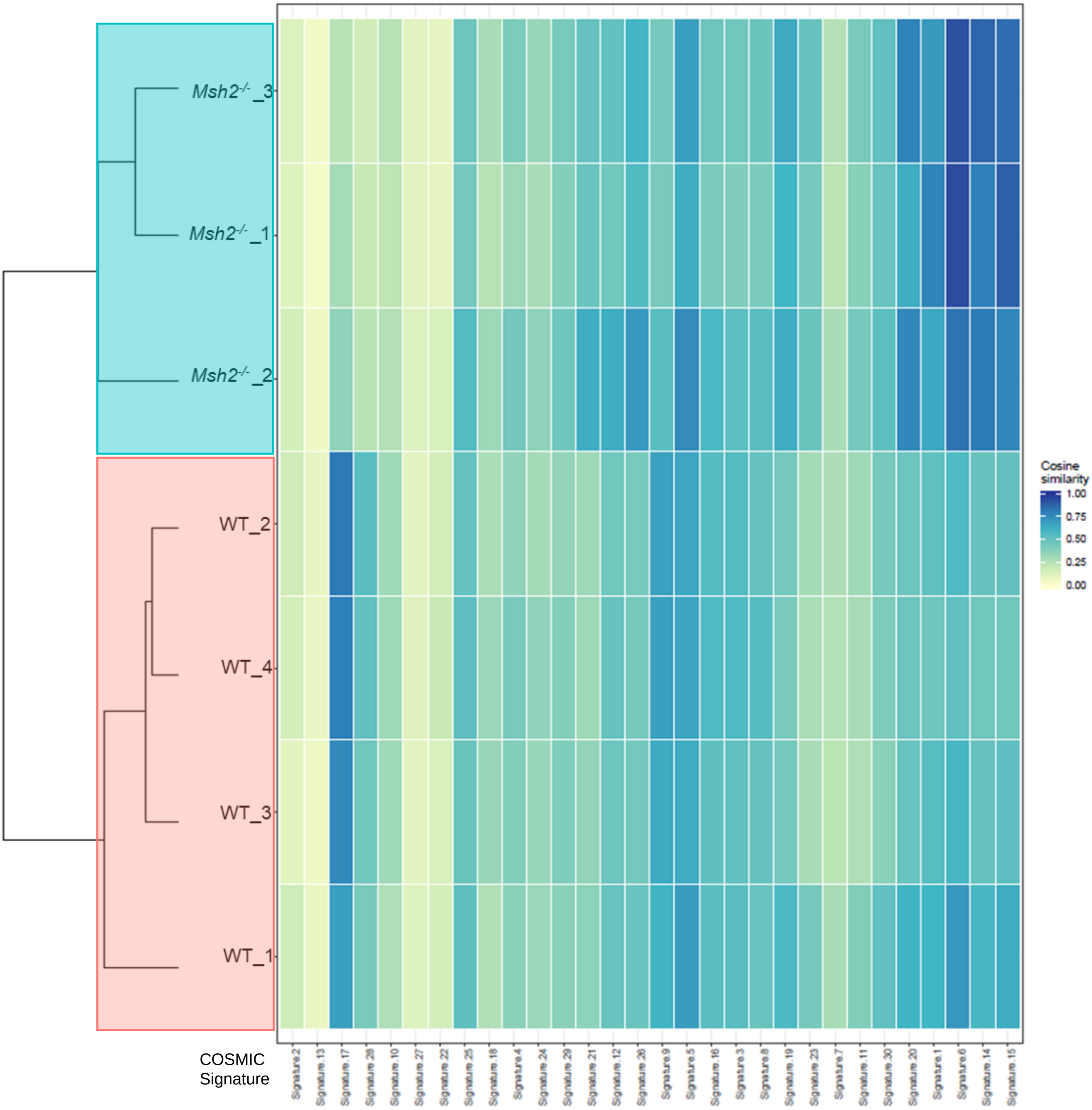


**Fig. S3. Cosine similarities between spectrum of each mouse and COSMIC signature.** Mice were clustered based on the similarities. Two distinct clusters composed of only the wild-type and *Msh2*$^{-/-}$ mice separately emerged as expected. The COSMIC signatures have been ordered according to hierarchical clustering.

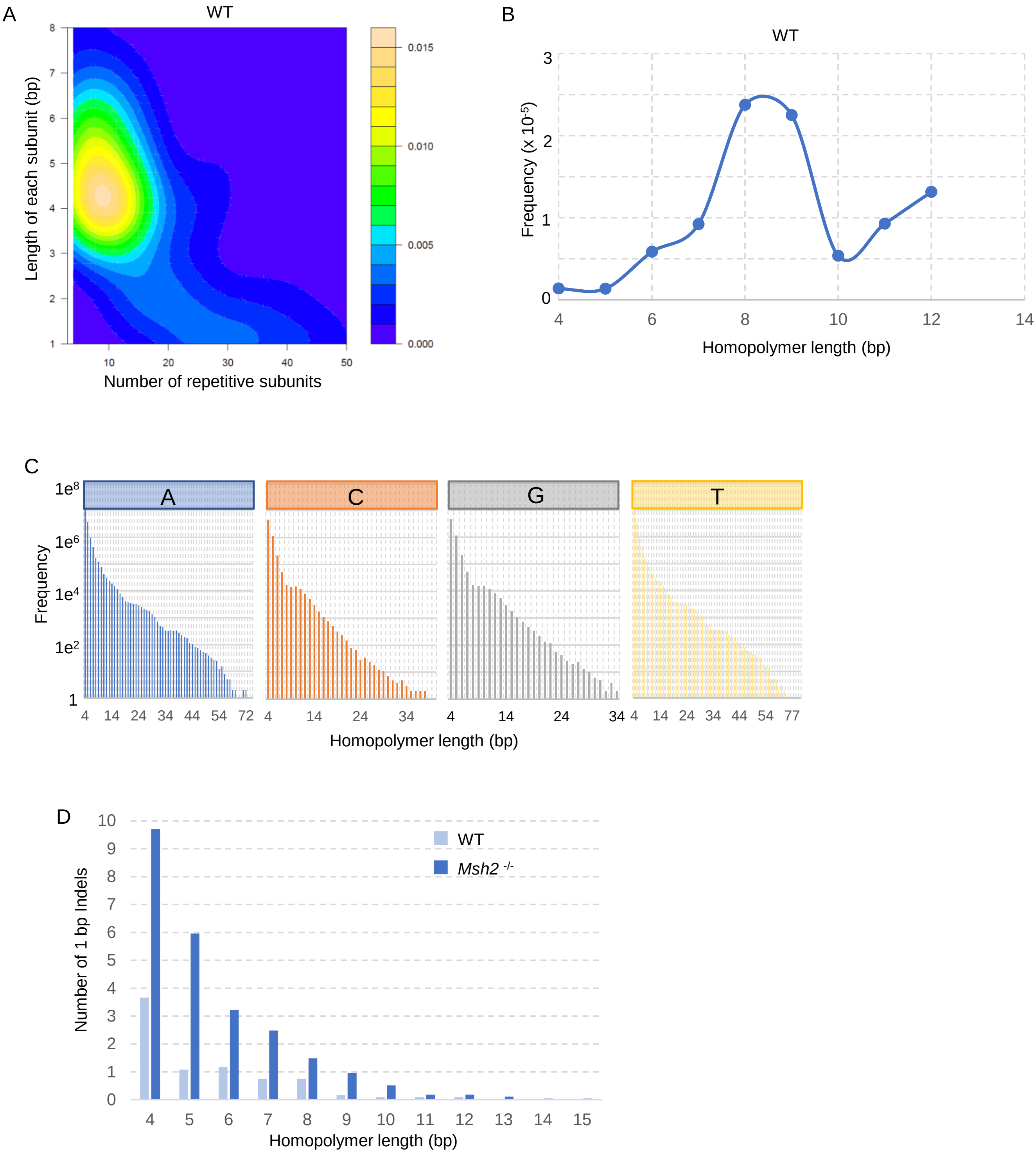


**Fig. S4. Distributions of 1-bp INDELs in simple repeats or homopolymers**. **A.** Distribution of 1-bp INDELs in simple repeats in wild-type mice. **B**. Frequency of 1-bp INDELs in homopolymers in wild-type mice. **C.** Frequency of homopolymers in the mouse reference genome. The Y-axis was shown in $log_{10}$ scale. **D.** Average number per cell of 1-bp INDELs in homopolymers in wild-type (12 cells, 4 mice) and *Msh2*$^{-/-}$ mice (9 cells, 3 mice).
